# Information theoretic evidence for predictive coding in the face processing system

**DOI:** 10.1101/089300

**Authors:** Alla Brodski-Guerniero, Georg-Friedrich Paasch, Patricia Wollstadt, Ipek Özdemir, Joseph T. Lizier, Michael Wibral

## Abstract

Predictive coding suggests that the brain infers the causes of its sensations by combining sensory evidence with internal predictions based on available prior knowledge. However, the neurophysiological correlates of (pre-)activated prior knowledge serving these predictions are still unknown. Based on the idea that such pre-activated prior knowledge must be maintained until needed we measured the amount of maintained information in neural signals via the active information storage (AIS) measure. AIS was calculated on whole-brain beamformer-reconstructed source time-courses from magnetoencephalography (MEG) recordings of 52 human subjects during the baseline of a Mooney face/house detection task. Pre-activation of prior knowledge for faces showed as alpha- and beta-band related AIS increases in content specific areas; these AIS increases were behaviourally relevant in brain area FFA. Further, AIS allowed decoding of the cued category on a trial-by-trial basis. Moreover, top-down transfer of predictions estimated by transfer entropy was associated with beta frequencies. Our results support accounts that activated prior knowledge and the corresponding predictions are signalled in low-frequency activity (<30 Hz).

**Significance statement:** Our perception is not only determined by the information our eyes/retina and other sensory organs receive from the outside world, but strongly depends also on information already present in our brains like prior knowledge about specific situations or objects. A currently popular theory in neuroscience, predictive coding theory, suggests that this prior knowledge is used by the brain to form internal predictions about upcoming sensory information. However, neurophysiological evidence for this hypothesis is rare – mostly because this kind of evidence requires making strong a-priori assumptions about the specific predictions the brain makes and the brain areas involved. Using a novel, assumption-free approach we find that face-related prior knowledge and the derived predictions are represented and transferred in low-frequency brain activity.

## Introduction

In the last decade, predictive coding theory has become a dominant paradigm to organize behavioral and neurophysiological findings into a coherent theory of brain function (George and Hawkins, 2009; Friston, 2010; Huang and Rao, 2011; Clark, 2012; Hohwy, 2013). Predictive coding theory proposes that the brain constantly makes inferences about the state of the outside world. This is supposed to be accomplished by building hierarchical internal predictions based on prior knowledge which are compared to incoming information in order to continuously adapt these internal models (Mumford, 1992; Rao et al., 1999; Friston, 2005, 2010)

The postulated use of predictions for inference requires several preparatory steps: First, task relevant prior knowledge passively stored in synaptic weights needs to be transferred into activated prior knowledge, i.e. information stored in neural activity (see Zipser et al., 1993 for a distinction of active/passive storage). Subsequently, (pre-)activated prior knowledge needs to be maintained until needed and transferred as a prediction in top-down direction to a lower cortical area, where it will be matched with incoming information (e.g. Mumford, 1992; Friston, 2005, 2010).

With respect to the neural correlates of activated prior knowledge and predictions we know that the prediction of specific features or object categories increases fMRI BOLD activity in the brain region at which the feature or category is usually processed (Puri et al., 2009; Esterman and Yantis, 2009; Kok et al., 2014). However, little is known about how the maintenance of pre-activated prior knowledge and the corresponding transfer of predictions are actually implemented in neural activity proper.

As a first step towards resolving this issue a microcircuit theory of predictive coding has been put forward, suggesting internal predictions to be processed in deep cortical layers and to manifest and to be transferred along descending fiber systems in low-frequency neural activity (<30 Hz) (Bastos et al., 2012).

This theory is in line with the findings of a spectral predominance of low-frequency neural activity in deep cortical layers (Buffalo et al., 2011) and the physiological findings linking feedback connections to alpha/beta frequency channels in monkeys (Bastos et al., 2015) and humans (Michalareas et al., 2016).

Recently, this microcircuit theory of predictive coding gained experimental support by neurophysiological studies showing the predictability of events to be associated with neural power in alpha (Bauer et al., 2014; Sedley et al., 2016) or beta frequencies (Pelt et al., 2016).

However, representation and signalling of pre-activated prior knowledge serving predictions has been difficult to investigate with classical analysis methods. One reason is that classical analysis methods require a-priori assumptions about which predictions specific brain areas are going to make - assumptions which might be very challenging to make beyond early sensory cortices and for complex experimental designs (Wibral et al., 2014, section 4.4, p. 9). Moreover, classical analysis methods do not allow quantifying the *amount* of pre-activated prior knowledge for predictions, as for instance diminished neural activity measured by fMRI, MEG/EEG may still come with less or more information being maintained in these signals. To overcome these problems we studied the maintenance and signalling of pre-activated prior knowledge for predictions using the information-theoretic measures of active information storage (AIS, see Methods in Lizier et al., 2012; also see Gómez et al., 2014 for an application to MEG), and transfer entropy (TE, Schreiber, 2000; Vicente et al., 2011a). AIS measures the amount of information in the future of a process predicted by its past (predictable information) while TE measures the amount of directed information transfer between two processes (see Methods for details).

Using these information-theoretic measures we investigated the pre-activation of prior knowledge for face predictions on neural source activity reconstructed from MEG recordings of 52 human subjects. In order to induce the pre-activation of face-related prior knowledge, subjects were instructed to detect Faces in two-tone stimuli (Mooney and Ferguson, 1951; Cavanagh, 1991).

## Methods

### Basic concept and testable hypotheses

To study the neural correlates of pre-activated prior knowledge for face predictions we used the information-theoretic measures active information storage (AIS) and transfer entropy (TE) - measuring predictable information (see Methods in Lizier et al., 2012) and information transfer (Schreiber, 2000; Vicente et al., 2011), respectively.

The use of AIS and TE in our study is based on the following rationale: Since the brain will usually not know exactly when a prediction will be needed, it will maintain activated prior knowledge related to the content of the prediction over time. If there is a reliable neural code that maps between content and activity, maintained activated prior knowledge must be represented as maintained information content in neural signals, measurable by AIS (Figure 1).

**Figure 1.**
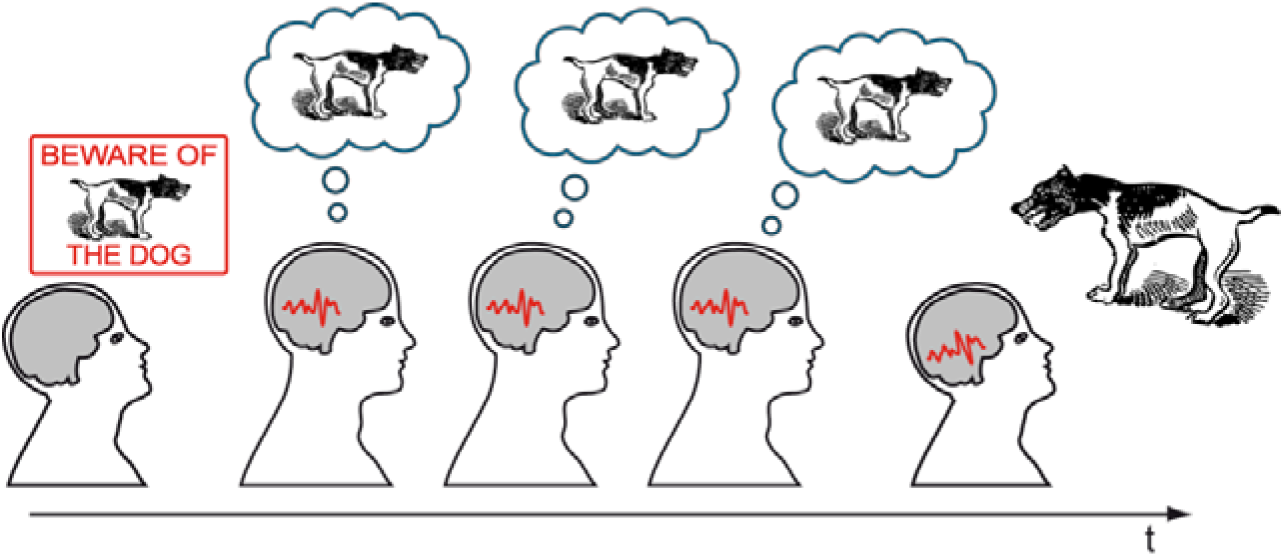
Central idea of the study. Typically, pre-activated prior knowledge related to the content of a prediction has to be maintained as the brain will not know exactly when it will be needed. If there is a reliable neural code that maps between content and activity, maintained activated prior knowledge should lead to brain signals that are themselves predictable over time (here the brain signals are depicted as identical, although the relation between past and future will almost certainly be much more complicated). Figure elements obtained from OpenCliparts Library (http://www.opencliparts.org) and modified.

Importantly, we do not suggest that predictable information in neural signals as measured by AIS measures the predictability of external events. Rather, we suggest that AIS can be used as a measure to detect increased predictable information in specific brain areas. This predictable information is bound to rise (see Fig. 1) when prior knowledge is pre-activated based on perceptual demands and thereby becomes available for predictions.

Further, predictions based on prior knowledge are supposed to be transferred to hierarchically lower brain areas, where they can be matched with incoming information. This information transfer thus must be measurable via TE.

From this basic concept we derived five testable hypotheses about AIS and TE in the predictive coding framework:

1. When activated prior knowledge is maintained, predictable information as measured by AIS is supposed to be high in brain areas specific to the content of the predictions.
2. If the microcircuit theory of predictive coding is correct, maintenance of pre-activated prior knowledge should be reflected in alpha/beta frequencies, i.e., predictable information and alpha/beta power should correlate.
3. If maintenance of relevant prior knowledge is reflected by predictable information on a trial-by-trial basis, the content of predictions should be also decodable from AIS information on a trial-by-trial basis.
4. Information transfer related to predictions (i.e. signalling of pre-activated prior knowledge measured by TE) should occur in a top-down direction from brain areas showing increased predictable information, and should be reflected in alpha/beta band Granger causality.
5. As predictions based on pre-activated prior knowledge are known to facilitate performance, predictable information is supposed to correlate with behavioural parameters, if it reflects the relevant pre-activated prior knowledge.

### Subjects

57 subjects participated in the MEG experiment. 5 of these subjects had to be excluded due to excessive movements, technical problems, or unavailability of anatomical scans. 52 subjects remained for the analysis (average age: 24.8 years, SD 2.8, 23 males). Each subject gave written informed consent before the beginning of the experiment and was paid 10€ per hour for participation. The local ethics committee (Johann Wolfgang Goethe University clinics, Frankfurt, Germany) approved of the experimental procedure. All subjects had normal or corrected-to-normal visual acuity and were right handed according to the Edinburgh Handedness Inventory scale (Oldfield, 1971). The large sample size subjects was chosen to reduce the risk of false positives, as suggested by (Button et al., 2013).

### Stimuli and stimulus presentation

Photographs of faces and houses were transformed into two-tone (black and white) images known as Mooney stimuli (Mooney and Ferguson, 1951). Mooney stimuli were used based on the rationale that recognition of two-tone stimuli cannot be accomplished without relying on prior knowledge from previous experience, as is evident for example from the late onset of two-tone image recognition capabilities during development (> 4 years of age, Mooney, 1957) and from theoretical considerations (Kemelmacher-Shlizerman et al., 2008).

In order to increase task difficulty, in addition to Mooney faces and houses also scrambled stimuli (SCR) were created from each of the resulting Mooney faces and Mooney houses by displacing the white or black patches within the given background. Thereby all low-level information was maintained but the configuration of the face or house was destroyed. Examples of the stimuli can be seen in Figure 2.

**Figure 2.**
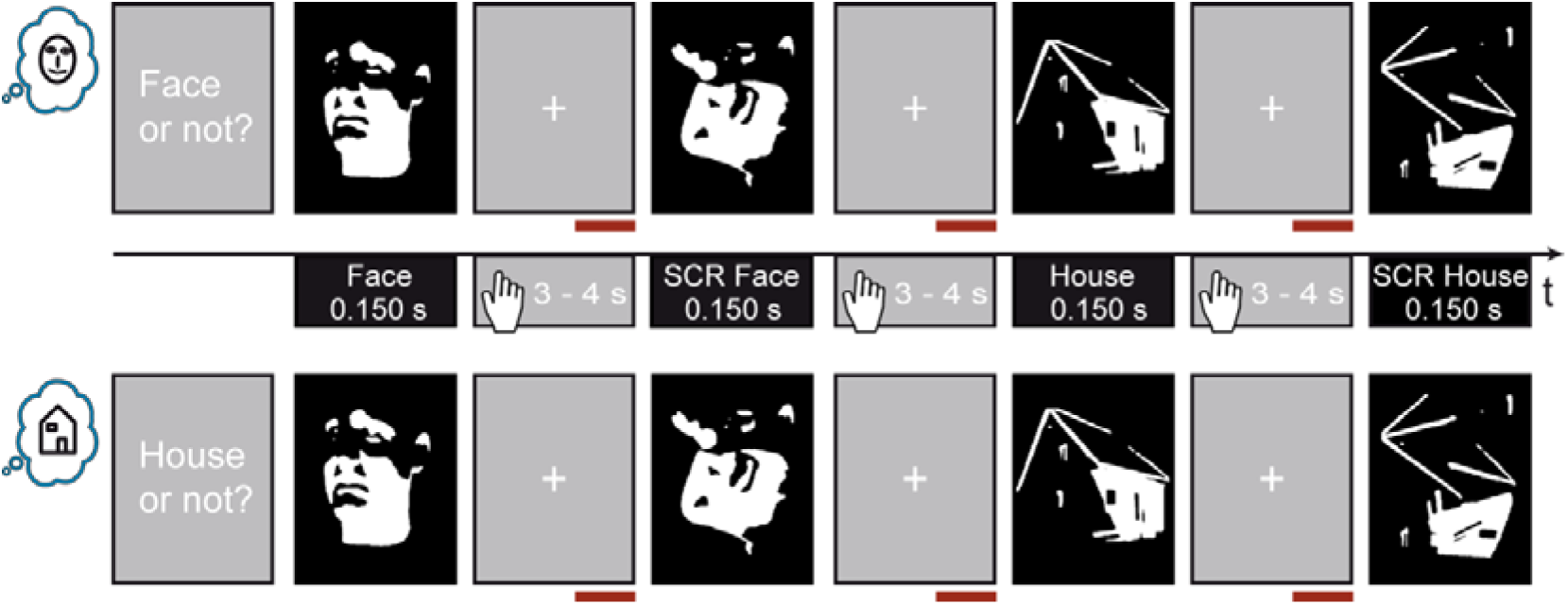
Experimental design. *Top* and *bottom:* Exemplary stimulus presentation in Face blocks *(top)* and in House blocks *(bottom).* Face and House icons on the left indicate Face and House blocks, respectively. *Middle:* Depiction of stimulus categories and timing. The beginning of the response time window is indicated by the hand icon. Red horizontal bars mark the analysis interval. SCR - Scrambled Mooney stimuli, not representing a face or house.

All stimuli were resized to a resolution of 591x754 pixels. Stimulus manipulations were performed with the program GIMP (GNU Image Manipulation Program, 2.4, free software foundation, Inc., Boston, Massachusetts, USA).

A projector with a refresh rate of 60 Hz (resolution 1024x768 pixels) was used to display the stimuli at the center of a translucent screen (background set to gray, 145 cd/m^2^). Stimulus presentation during the experiment was controlled using the Presentation software package (Version 9.90, Neurobehavioral Systems).

The experiment consisted of eight blocks of seven minutes. In each block 120 stimuli were presented (30 Mooney faces, 30 Mooney houses, 30 SCR faces, 30 SCR houses) in a randomized order. Stimuli were presented for 150 ms with a vertical visual angle of 24.1 and a horizontal visual angle of 18.8 degrees. The inter-trial-interval between stimulus presentations was randomly jittered from 3 to 4 seconds (in steps of 100 ms).

### Task and Instructions

Subjects performed a detection task for faces or houses (Figure 2). Each of the eight experimental blocks started with the presentation of a written instruction; four of the experimental blocks started with the instruction “Face or not?” while for the other four experimental blocks started with the instruction “House or not?”. The former are referred to as “Face blocks” and the latter as “House blocks”. Face and House blocks were presented in alternating order. The same blocks of stimuli were presented as Face blocks for half of the subjects, while for the other half of the subjects these experimental blocks appeared as House blocks and vice versa. This way, the initial block was alternated between subjects (i.e. half of the subjects started with Face blocks and the other half with House blocks). Importantly, as the blocks contained the same face, house, SCR face and SCR house stimuli the only difference between face and house blocks was in the subjects’ instruction.

To avoid accidental serial effects, the order of blocks was reversed for half of the subjects. Subject responded by pressing one of two buttons directly after stimulus presentation. The button assignment for a ‘Face’ or ‘No-Face’ response in Face blocks and ‘House’ or ‘No-House’ block was counterbalanced across subjects (n=26 right index finger for ‘Face’ response).

Between stimulus presentations, subjects were instructed to fixate a white cross on the center of the gray screen. Further, they were instructed to maintain fixation during the whole block and to avoid any movement during the acquisition session. Before data acquisition, subjects performed Face and House test blocks of two minutes with stimuli not used during the actual task. During the test blocks subjects received a feedback whether their response was correct or not. No feedback was provided during the actual task.

### Data acquisition

MEG data acquisition was performed in line with recently published guidelines for MEG recordings (Gross et al., 2012). MEG signals were recorded using a whole-head system (Omega 2005; VSM MedTech Ltd.) with 275 channels. The signals were recorded continuously at a sampling rate of 1200 Hz in a synthetic third-order gradiometer configuration and were filtered online with fourth-order Butterworth filters with 300 Hz low pass and 0.1 Hz high pass.

Subjects’ head position relative to the gradiometer array was recorded continuously using three localization coils, one at the nasion and the other two located 1 cm anterior to the left and right tragus on the nasion-tragus plane for 43 of the subjects and at the left and right ear canal for 9 of the subjects.

For artefact detection the horizontal and vertical electrooculogram (EOG) was recorded via four electrodes; two were placed distal to the outer canthi of the left and right eye (horizontal eye movements) and the other two were placed above and below the right eye (vertical eye movements and blinks). In addition, an electrocardiogram (ECG) was recorded with two electrodes placed at the left and right collar bones of the subject. The impedance of each electrode was kept below 15 kΩ.

Structural magnetic resonance (MR) images were obtained with either a 3T Siemens Allegra or a Trio scanner (Siemens Medical Solutions, Erlangen, Germany) using a standard T1 sequence (3-D magnetization-prepared-rapid-acquisition gradient echo sequence, 176 slices, 1 x 1 x 1 mm voxel size). For the structural scans vitamin E pills were placed at the former positions of the MEG localization coils for co-registration of MEG data and magnetic resonance images.

Behavioral responses were recorded using a fiberoptic response pad (Photon Control Inc. Lumitouch Control ™ Response System) in combination with the Presentation software (Version 9.90, Neurobehavioral Systems).

### Statistical analysis of behavioral data

Responses were classified as correct or incorrect based on the subject’s first answer. For hit rate analysis the accuracy for each condition was calculated. For reaction time analysis only correct responses were considered.

Post-hoc Wilcoxon signed rank tests were performed on hitrates as well as reaction times. To account for multiple testing, sequential Bonferroni-Holm correction (Holm, 1979) was applied (uncorrected alpha = 0.05).

### MEG-data preprocessing

MEG Data analysis was performed with Matlab (RRID:nlx_153890; Matlab 2012b, The Mathworks, Inc.) using the open source Matlab toolbox Fieldtrip (RRID:nlx_143928; Oostenveld et al., 2011); Version 2013 11-11) and custom Matlab scripts.

Only trials with correct behavioral responses were taken into account for MEG data analysis. The focus of data analysis was on the prestimulus intervals from 1 s to 0.050 s before stimulus onset. Trials containing sensor jump-, or muscle-artefacts were rejected using automatic FieldTrip artefact rejection routines. Line noise was removed using a discrete Fourier transform filter at 50,100 and 150 Hz. In addition, independent component analysis (ICA; (Makeig et al., 1996) was performed using the extended infomax (runica) algorithm implemented in fieldtrip/EEGLAB. ICA components strongly correlated with EOG and ECG channels were removed from the data. Finally, data was visually inspected for residual artefacts.

In order to minimize movement related errors, the mean head position over all experimental blocks was determined for each subject. Only trials in which the head position did not deviate more than 5 mm from the mean head position were considered for further analysis.

As artefact rejection and trial rejection based on the head position may result in different trial numbers for Face and House blocks, after trial rejection the minimum amount of trials across Face and House blocks was selected randomly from the available trials in each block (stratification).

### Sensor level spectral analysis

Spectral analysis at the sensor level was performed in order to determine the subdivision of the power spectrum in frequency bands (see Brodski et al., 2015 for a similar approach). As we aimed to identify frequency bands based on stimulus related increases or decreases, respectively, before spectral analysis new data segments were cut from −0.55 to 0.55 s around stimulus onset. For spectral analysis we used a multitaper approach (Percival and Walden, 1993) based on Slepian sequences (Slepian, 1978). The spectral transformation was applied in an interval from 4 to 150 Hz in 2 Hz steps in time steps of 0.01 s and using two slepian tapers for each frequency. For each subject, time-frequency representations were averaged for Face blocks and House blocks as well as within the time interval of “baseline” (-0.35 s – 0.05 s) and “task” (0.05 s – 0.35 s), respectively. Average spectra of task and baseline period were contrasted over subjects using a dependent-sample permutation t-metric with a cluster based correction method (Maris and Oostenveld, 2007) to account for multiple comparisons. Adjacent samples whose *t*-values exceeded a threshold corresponding to an uncorrected α-level of 0.05 were defined as clusters. The resulting cluster sizes were then tested against the distribution of cluster sizes obtained from 1000 permuted datasets (i.e. labels “task” and “baseline” were randomly reassigned within each of the subjects). Cluster sizes larger than the 95^th^ percentile of the cluster sizes in the permuted datasets were defined as significant.

Following the same approach as (Brodski et al., 2015) based on the significant clusters of the task vs. baseline statistics five frequency bands were defined for further analysis: (1) 8–14 Hz (alpha); (2) 14–32 Hz (beta); (3) 32–56 Hz (low gamma); (4) 56–64 Hz (mid gamma) and (5) 64–150 Hz (high gamma) (Figure 3).

**Figure 3.**
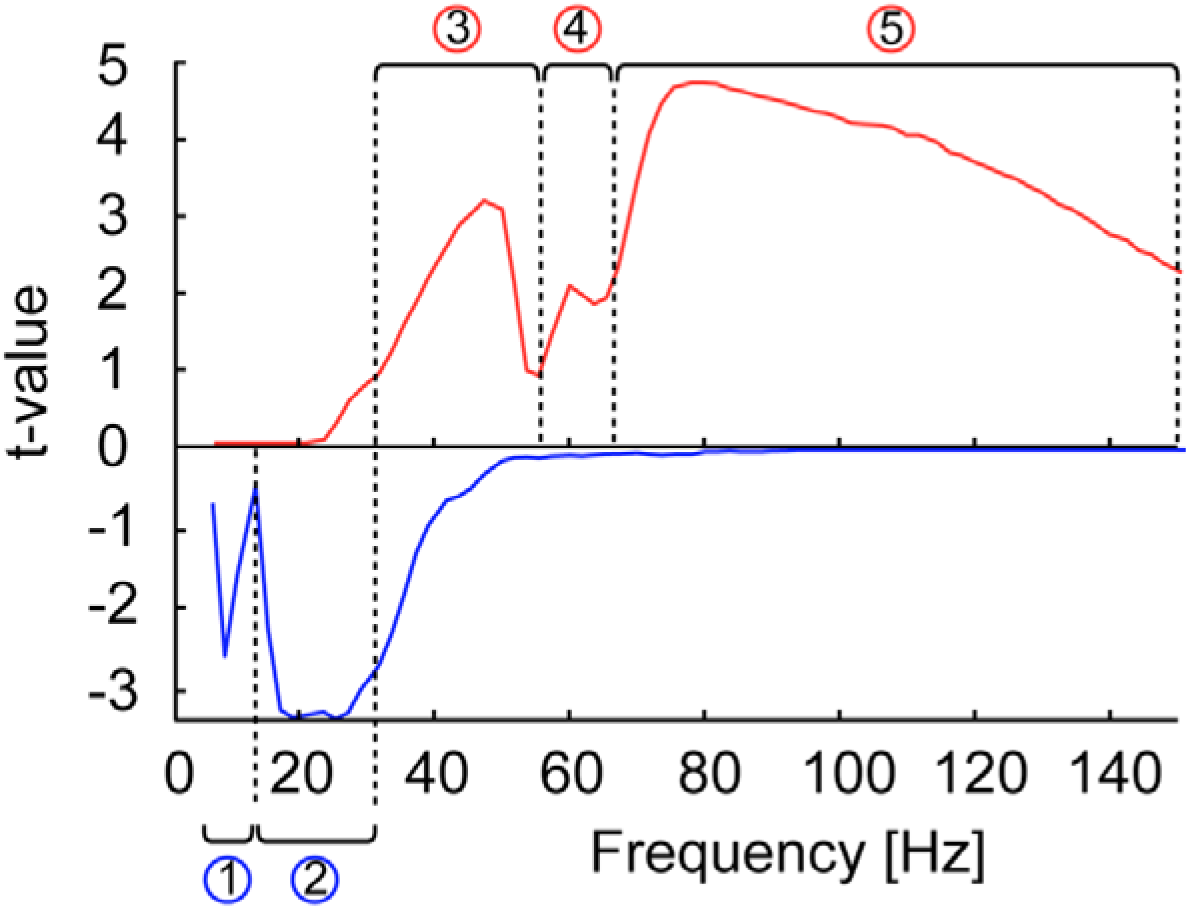
Sensor-level frequency analysis - defining frequency bands. Power spectra for all of the significant clusters (one positive and one negative cluster) at the sensor level (permutation t-metric, contrast [0.05s 0.35s] vs [-0.35s −0.05s] around stimulus onset, *t* values masked by p < 0.05, cluster correction, n = 52). Frequency analysis at the sensor level was calculated using both blocks types jointly. Task-related increases in power are shown in red (positive cluster) and task-related decreases in blue (negative cluster). Black dashed lines frame the identified frequency ranges.

### Source grid creation

In order to create individual source grids we transformed the anatomical MR images to a standard T1 MNI template from the SPM8 toolbox (http://www.fil.ion.ucl.ac.uk/spm) - obtaining an individual transformation matrix for each subject. We then warped a regular 3-D dipole grid based on the standard T1 template (spacing 15mm resulting in 478 grid locations) with the inverse of each subjects’ transformation matrix, to obtain an individual dipole grid for each subject in subject space. This way, each specific grid point was located at the same brain area for each subject, which allowed us to perform source analysis with individual head models as well as multi-subject statistics for all grid locations. Lead-fields at those grid locations were computed for the individual subjects with a realistic single shell forward model (Nolte, 2003) taking into account the effects of the ICA component removal in pre-processing.

### Source time course reconstruction

To enable a whole brain analysis of active information storage (AIS), we reconstructed the source time courses for all 478 source grid locations.

For source time course reconstruction we calculated a time-domain beamformer filter (linear constrained minimum variance, LCMV; Van Veen et al., 1997) based on broadband filtered data (8 Hz high pass, 150 Hz low pass) from the prestimulus interval (−1 s to −0.050 s) of Face blocks as well as House blocks (use of common filters - see Gross et al., 2012, page 357).

For each source location three orthogonal filters were computed (x, y, z direction). To obtain the source time courses, the broadly filtered raw data was projected through the LCMV filters resulting in three time courses per location. On these source time courses we performed a singular value decomposition to obtain the time course in direction of the dominant dipole orientation. The source time course in direction of the dominant dipole orientation was used for calculation of active information storage (AIS).

### Definition of active information storage

We assume that the reconstructed source time courses for each brain location can be treated as realizations {*x*_1_,…, *x_t_*, …, *x_N_*} of a random process *X =* {*X*_1_,…, *X_t_*, …, *X_N_*}, which consists of a collection of random variables, *X_t_*, ordered by some integer *t*. AIS then describes how much of the information the next time step *t* of the process is predictable from its immediate past state (Lizier et al., 2012). This is defined as the mutual information

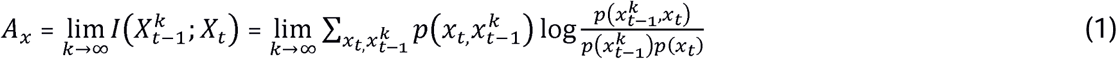

where *I* is the mutual information and *p*(.) are the variables' probability density functions. Variable 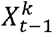 describes the past *state* of *X* as a collection of past random variables 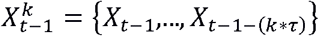, where *k* is the embedding dimension, i.e., the number of time steps used in the collection, and τ the embedding delay between these time steps. For practical purposes, *k* has to be set to a finite value *k_max_*, such that the history before time point *t* – *k_max_* * *τ* does (statistically) not further improve the prediction of *X_t_* from its past (Lizier et al., 2012).

Predictable information as measured by AIS indicates that a signal is both rich in information and predictable at the same time. Note that neither a constant signal (predictable but low information content) nor a memory-less stochastic process (high information content but unpredictable) will exhibit high AIS values. In other words, a neural process with high AIS must visit many different possible states (rich dynamics), yet visit these states in a predictable manner with minimal branching of its trajectory (this is the meaning of the log ratio of equation (1)). As such, AIS is a general measure of information that is maintained in a process, and could here reflect any form of memory based on neural activity. AIS is linked specifically to activated prior knowledge in our study via the experimental manipulation that alternately activates face- or house-specific prior knowledge, and by investigating the difference in AIS between the two conditions..

### Analysis of predictable information using active information storage

The history dimension (*k*_max_; range 3 to 6) and optimal embedding delay parameter (tau; range 0.2 to 0.5 in units of the autocorrelation decay time) was determined for each source location separately using Ragwitz’ criterion (Ragwitz and Kantz, 2002), as implemented in the TRENTOOL toolbox (Lindner et al., 2011). To avoid a bias in estimated values based on different history dimensions, we chose the maximal history dimension across Face and House blocks for each source location (median *k*_max_ over source locations and subjects =4).

The actual spacing between the time-points in the history was the median across trials of the output of Ragwitz’ criterion for the embedding delay tau (Lindner et al., 2011).

Based on the assumption of stationarity in the prestimulus interval, AIS was computed on the embedded data across all available time points and trials. This was done separately for each source location and condition in every subject.

Computation of AIS was performed using the Java Information Dynamics Toolkit (Lizier, 2014). A minimum of 68400 samples entered the AIS analysis for each subject, block type and source location (minimum of 57 trials, approx. 1 sec time interval, sampling rate 1200 Hz). AIS was estimated with 4 nearest neighbours in the joint embedding space using the Kraskov-Stoegbauer-Grassberger (KSG) estimator (Kraskov et al., 2004); algorithm 1), as implemented in the open source Java Information Dynamics Toolkit (JIDT; Lizier, 2014))

Computation of AIS was performed at the Center for Scientific Computing (CSC) Frankfurt, using the high-performance computing Cluster FUCHS (https://csc.uni-frankfurt.de/index.php?id=4), which enabled the computationally demanding calculation of AIS for the whole brain across all subjects as well as Face and House blocks (478 x 52 x 2 = 49712 computations of AIS).

### AIS Statistics

In order to determine the source locations in which AIS values were increased when subjects held face information in memory, a within-subject permutation t-metric was computed. Here, AIS values for each source location across all subjects were contrasted for Face blocks and House blocks. The permutation test was chosen as the distribution of AIS values is unknown and not assumed to be Gaussian. To account for multiple comparisons across the 478 source locations, a cluster-based correction method (Maris and Oostenveld, 2007) was used. Clusters were defined as adjacent voxels whose t-values exceeded a critical threshold corresponding to an uncorrected alpha level of 0.01. In the randomization procedure labels of Face block and House block data were randomly reassigned within each subject. Cluster sizes were tested against the distribution of cluster sizes obtained from 5000 permuted data sets. Cluster values larger than the 95^th^ percentile of the distribution of cluster sizes obtained for the permuted data sets were considered to be significant.

## Correlation analysis

We investigated the relationship of spectral power in the prestimulus interval and AIS values on the single trial level. Before calculation of the correlation coefficient, single trial spectral power in each of the predefined frequency bands and single trial AIS values were z-normalized for each subject. These values were appended for Face and House blocks, pooled over all subjects and Spearman’s rho was calculated. Then, trials were shuffled 1000 times for spectral power and AIS values separately within each subject and correlation analysis was repeated for each randomization. Original correlation values larger than the 95^th^ percentile of the distribution of correlation values in the shuffled data were considered as significant. This statistical procedure conforms to a permutation test of the correlation where permutations are restricted within the levels of the factor subjects.

We also calculated the correlation of t-values computed from AIS (based on the dependent sample t-metric, contrast Face blocks vs. House blocks) for all grid points at the source level with the t-values obtained from the same contrast based on beamformer reconstructed source power in the alpha (8-14 Hz) and beta (14-32 Hz) frequency band. Source power was reconstructed with the DICS (dynamic imaging of coherent sources, Gross et al., 2001) algorithm as implemented in the FieldTrip toolbox using real values filter coefficients only - see also Grützner et al., 2010).

Last, we accessed the relationship of AIS values and reaction times for each subject. To this end before the correlation analysis for each subject mean reaction times and mean AIS values in the brain areas of interest for Face and House blocks were subtracted from each other. This allowed accounting for differential behavioral speed between subjects. The correlation of the difference in AIS values and the difference in reaction times was calculated via Spearman skipped correlations using the Robust correlation Toolbox (Pernet et al., 2013). Calculation of skipped correlations includes identifying and removing bivariate outliers (Rousseeuw, 1984; Rousseeuw and Driessen, 1999; Verboven and Hubert, 2005). This can provide a more robust measure, which has been recommended for brain-behaviour correlation analyses (Rousselet and Pernet, 2012). The uncorrected alpha level was set to 0.05. For each correlation bootstrap confidence intervals (CIs) were computed based on 1000 resamples. In order to account for multiple comparisons across brain areas, bootstrap CIs were adjusted using Bonferroni correction. If the adjusted CI did not encompassed 0, the correlation was considered as significant.

## Decoding analysis

To investigate whether prediction content (i.e. face or house block) can be decoded from individual trial AIS values, we applied a multivariate analysis using support vector machines (SVMs) with the libsvm toolbox (Chang and Lin, 2011; available at http://www.csie.ntu.edu.tw/~cjlin/libsvm). For each subjects the linear SVM classifier was trained using 70% randomly chosen trials as training data. However, the training data contained always the same amount of trials for face and house blocks, respectively. Parameters for the SVMs were optimized in a three-fold cross-validation procedure for the training data only. Subsequently, the classifier was tested using the data from the remaining 30% of the trials with the best parameters obtained from the training procedure, thereby ensuring strict separation of training and testing data (Nowotny, 2014).

This procedure was repeated 10 times. We report the median accuracy value for each subject. In order to test the significance of the median accuracy value, for each subject the labels of face blocks and house blocks were randomly permuted 500 times for each of the 10 training and testing sets and the median over the 10 accuracy values was calculated also for the permuted data sets. A median accuracy value larger than the 99.999% (threshold Bonferroni adjusted for the 52 multiple comparisons) of the permuted median accuracy values obtained for the permuted data sets was considered to be significant, corresponding to an un-corrected alpha level of 0.05.

## Definition of transfer entropy (and Granger analysis)

Transfer entropy (TE, (Schreiber, 2000) was applied to investigate the information transfer between the brain areas identified with AIS analysis. For links with significant information transfer, we post-hoc studied the spectral fingerprints of these links using spectral Granger analysis (Granger, 1969).

Both, TE and Granger analysis are implementations of Wieners principle (Wiener, 1956)which in short can be rephrased as follows: If the prediction of the future of one time series X, can be improved in comparison to predicting it from the past of X alone by adding information from the past of another time series Y, then information is transferred from Y to X.

TE is an information-theoretic, model-free implementation of Wiener's principle and can be used, in contrast to Granger analysis, in order to study linear as well as non-linear interactions (e.g. Chang and Lin, 2011) and was previously applied to broadband MEG source data (Wibral et al., 2011). TE is defined as a conditional mutual information

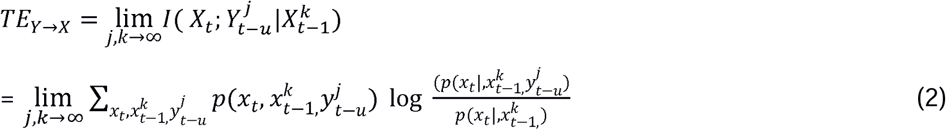

where *X_t_* describes the future of the target time series *X*, 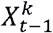 describes the past state of *X*, and 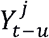 describes the past state of the source time series *Y*. As for the calculation of AIS, past states are defined as collections of past random variables with number of time steps *j* and *k* and a delay τ. The parameter *u* accounts for a physical delay between processes *Y* and *X* (Wibral et al., 2013) and can be optimized by finding the maximum TE over a range of assumed values for *u.*

## Analysis of information transfer using transfer entropy and Granger causality analysis

We performed TE analysis with the open-source Matlab toolbox TRENTOOL (Lindner et al., 2011), which implements the KSG-estimator (Kraskov et al., 2004; Frenzel and Pompe, 2007; Gómez-Herrero et al., 2015) for TE estimation. We used ensemble estimation (Wollstadt et al., 2014; Gómez-Herrero et al., 2015), which estimates TE from data pooled over trials to obtain more data and hence more robust TE-estimates. Additionally, we used Faes’ correction method to account for volume conduction (Faes et al., 2013).

In the TE analysis the same time intervals (prestimulus) and embedding parameters as for AIS analysis were used. TE values for Face blocks and House blocks were contrasted using a dependent-sample permutation t-metric for statistical analysis across subjects. In the statistical analysis, FDR correction was used to account for multiple comparisons across links (uncorrected alpha level 0.05). As for AIS, the history dimension for the past states was set to finite values; we here set *j*_max_ = *k*_max_ and used the values obtained during AIS estimation for the target time series of each signal combination.

For the significant TE links post-hoc nonparametric bivariate Granger causality analysis in the frequency domain (Dhamala et al., 2008) was computed. Using the nonparametric variant of Granger causality analysis avoids choosing an autoregressive model order, which may easily introduce a bias. In the nonparametric approach Granger causality is computed from a factorization of the spectral density matrix, which is based on the direct Fourier transform of the time series data (Dhamala et al., 2008). The Wilson algorithm was used for factorization (Wilson, 1972). A spectral resolution of 2 Hz and a spectral smoothing of 5 Hz were used for spectral transformation using the multitaper approach (Percival and Walden, 1993) (9 Slepian tapers). We were interested in the differences of Granger spectral fingerprints of Face and House blocks, however we also wanted to make sure that the Granger values for these difference significantly differed from noise. For that reason we created two additional “random” conditions by permuting the trials for the Face block and the House block condition for each source separately. Two types of statistical comparisons were performed for the frequency range between 8 and 150 Hz and each of the significant TE links: 1. Granger values in Face blocks were contrasted with Granger values in House blocks using a dependent-samples permutation t-metric 2. Granger values in Face blocks/House blocks were contrasted with the random Face block condition / random House block condition using another dependent-samples permutation t-metric. For the first test a cluster-correction was used to account for multiple comparisons across frequency (Maris and Oostenveld, 2007). Adjacent samples which uncorrected p-values were below 0.01 were considered as clusters. 5000 permutations were performed and the alpha value was set to 0.05. Frequency intervals in the Face block vs. House block comparison were only considered as significant if all included frequencies also reached significance in the comparison with the random conditions using a Bonferroni-Holms correction to account for multiple comparisons.

## Results

### Behavioral results

We found no differences between Face blocks and House blocks for hitrates (avg. hitrate Face blocks 93.9%; avg. hitrate House blocks 94.6%; Wilcoxon Signed rank test p=0.57) and reaction times of correct responses (avg. mean reaction times Face blocks 0.545 s, avg. reaction times House blocks 0.546 s; Wilcoxon Signed rank test p=0.85). Subjects showed equivalent behavioural patterns for both block types, for instance increased reaction times for the instructed stimulus conditions as these stimuli had to be distinguished from a similar distractor (SCR stimuli) (see Figure 4 for the analysis of behavioural differences between stimulus conditions within block types).

**Figure 4.**
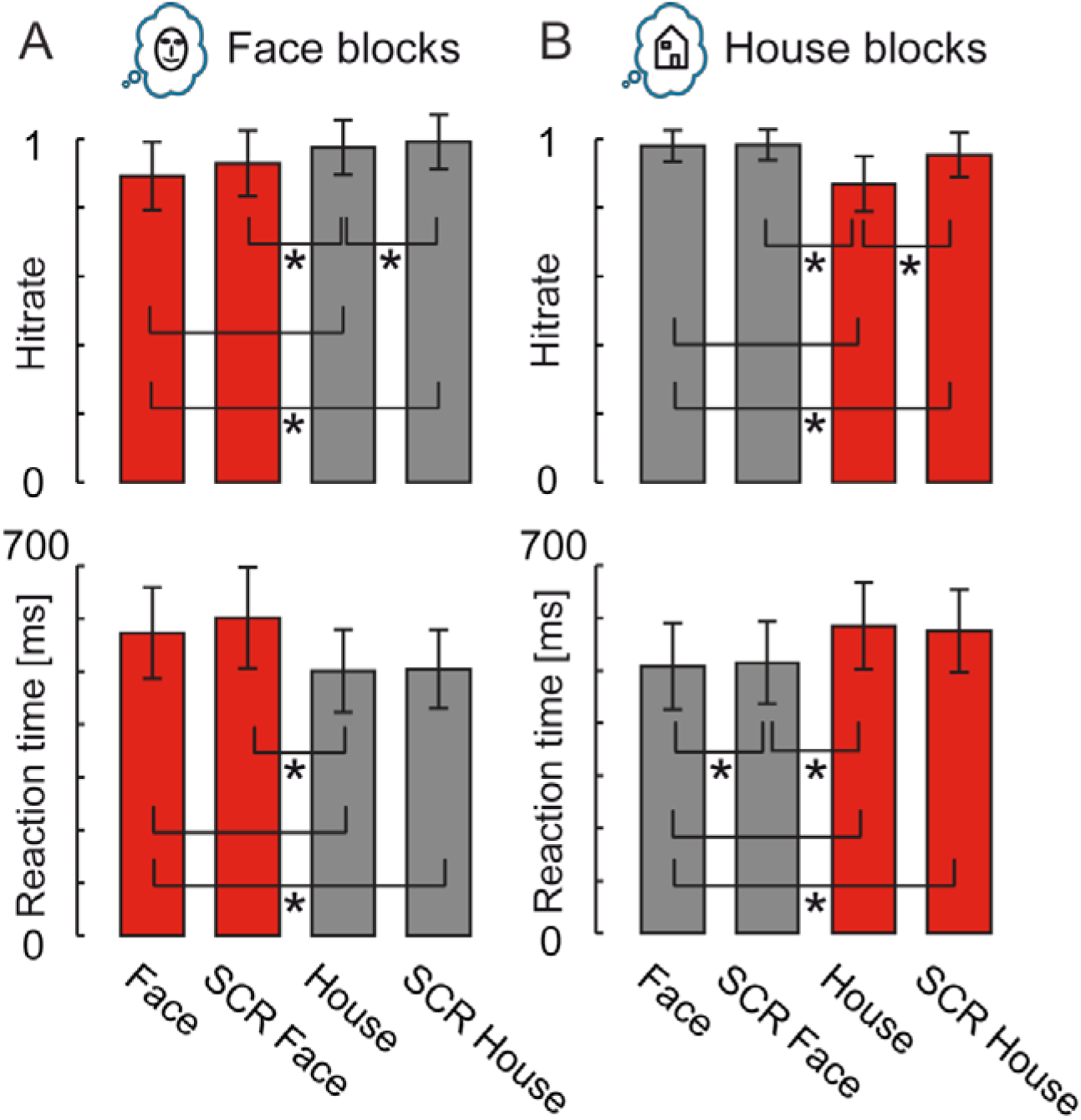
Behavioral results. Depiction of hitrates and reaction times of correct responses for (A) Face blocks and (B) House blocks. Equivalent conditions in different block types are marked in red and grey, respectively. Asterisks indicate significant differences based on Wilcoxon signed-rank tests within block type (n = 52; Bonferroni-Holms corrected for multiple comparisons). Error bars indicate standard deviation. SCR – scrambled Mooney stimuli.

### Analysis of predictable information

Statistical comparisons of AIS values between Face blocks and House blocks in the prestimulus interval revealed increased AIS values for Face blocks in clusters in fusiform face area (FFA), anterior inferior temporal cortex (aIT), occipital face area (OFA), posterior parietal cortex (PPC) and primary visual cortex (V1) (Figure 5). We referred to these five brain areas as “face prediction network” and subjected it to further analyses. In contrast to this finding of a face prediction network, we did not find brain areas showing significantly higher AIS values in House blocks compared to Face blocks. This is similar to highly cited previous studies that failed to find prediction effects for houses in the brain in contrast to faces (e.g. Summerfield et al., 2006a, 2006b; Trapp et al., 2015).

**Figure 5.**
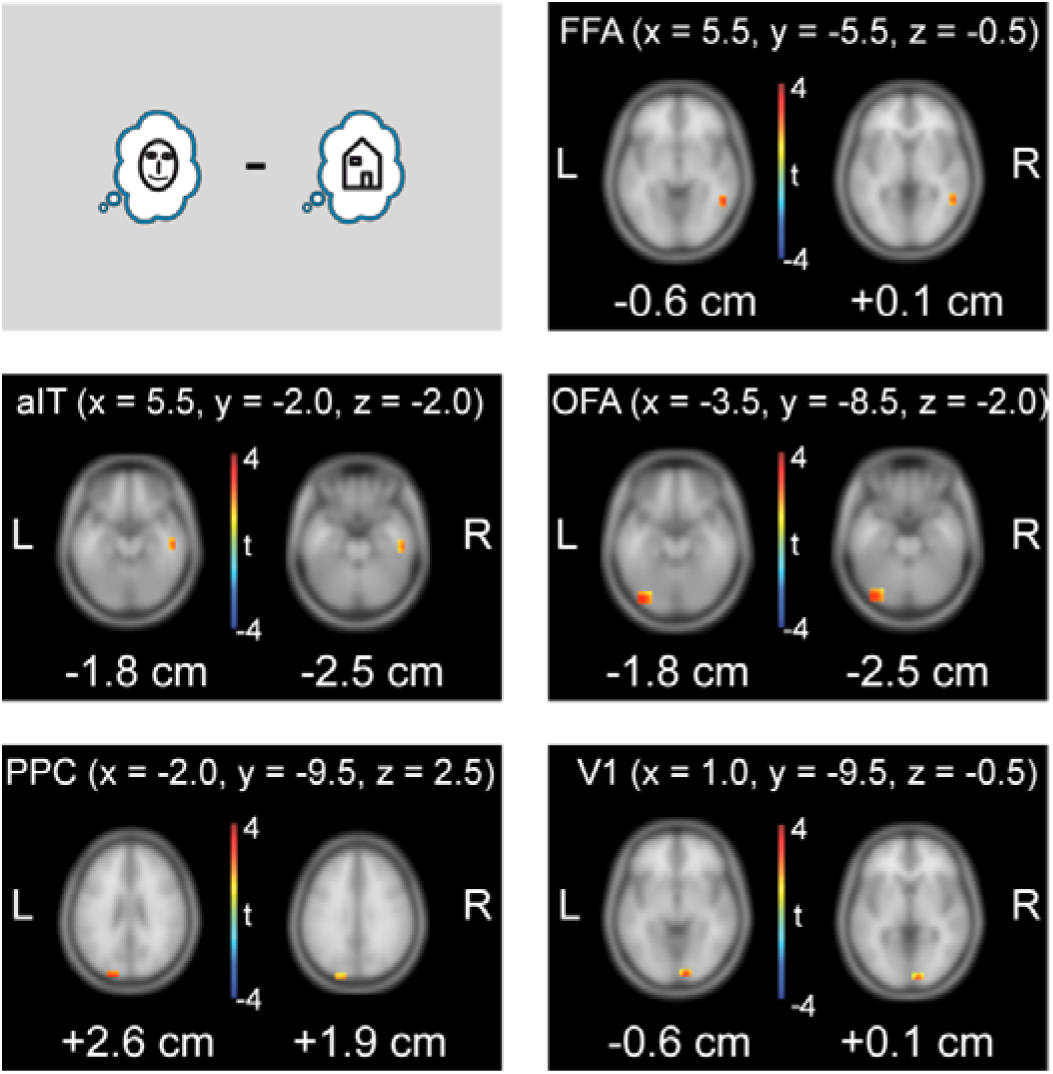
Statistical analysis of predictable information (measured by AIS) at the MEG source level. Results of whole-brain dependent samples permutation t-metric contrasting Face blocks and House blocks (n=52, t-values masked by p<0.05, cluster correction). Peak voxel coordinates in MNI space are shown at the top for each brain location; z-values are displayed below each brain slice. OFA = occipital face area; FFA = fusiform face area; aIT= anterior inferior temporal cortex; PPC = posterior parietal cortex; V1 = primary visual cortex

### Correlation of single trial power and single trial predictable information

In order to investigate the neurophysiological correlates of activated prior knowledge identified via AIS analysis, a correlation analysis of single trial power in distinct frequency bands with single trial AIS was conducted. Correlation analysis revealed a strong positive correlation in the alpha and beta frequency bands, only very small mostly positive correlations in the low and mid-gamma frequency bands and a small negative correlation for the high-gamma frequency band (Table 1). Note that although the correlations in the higher frequency bands were partly significant, the effect size was much higher in the alpha and beta frequency band. This means that alpha and beta band activity is the most likely carrier of activated prior knowledge.

**Table 1:** Correlation of single trial power and single trial predictable information (measured by AIS) in the face prediction network

|  | OFA | aIT | PPC | V1 | FFA |
| --- | --- | --- | --- | --- | --- |
| 8-14 Hz<br>(alpha) | $\rho = 0.58$<br>$p < 0.001$ | $\rho = 0.54$<br>$p < 0.001$ | $\rho = 0.59$<br>$p < 0.001$ | $\rho = 0.61$<br>$p < 0.001$ | $\rho = 0.58$<br>$p < 0.001$ |
| 14-32 Hz<br>(beta) | $\rho = 0.53$<br>$p < 0.001$ | $\rho = 0.52$<br>$p < 0.001$ | $\rho = 0.53$<br>$p < 0.001$ | $\rho = 0.52$<br>$p < 0.001$ | $\rho = 0.54$<br>$p < 0.001$ |
| 32-56 Hz<br>(low gamma) | $\rho = 0.13$<br>$p < 0.001$ | $\rho = 0.10$<br>$p < 0.001$ | $\rho = 0.17$<br>$p < 0.001$ | $\rho = 0.14$<br>$p < 0.001$ | $\rho = 0.13$<br>$p < 0.001$ |
| 56-64 Hz<br>(Mid-gamma) | $\rho = 0.06$<br>$p < 0.001$ | $\rho = -0.004$<br>$p = 0.22$ | $\rho = 0.07$<br>$p < 0.001$ | $\rho = 0.07$<br>$p < 0.001$ | $\rho = 0.02$<br>$p < 0.001$ |
| 64-150 Hz<br>(High-gamma) | $\rho = -0.07$<br>$p < 0.001$ | $\rho = -0.14$<br>$p < 0.001$ | $\rho = -0.04$<br>$p < 0.001$ | $\rho = -0.04$<br>$p < 0.001$ | $\rho = -0.09$<br>$p < 0.001$ |

While we found a significant correlation of single trial power and predictable information in the alpha and beta band, the contrast map over all source grid points for Face and House blocks (t-values obtained from dependent sample t-metric over subjects) did not correlate with the AIS contrast map for both, alpha and beta power (alpha rho = 0.043, p = 0.33; beta rho = 0.05, p =0.21). This suggests that AIS analysis provides additional information not directly provided by a spectral analysis. In sum, while AIS seems to be carried by alpha/beta-band activity, not all alpha/beta band activity contributes to AIS.

### Decoding prediction content from single trial AIS values

To study whether face or house predictions can be decoded from AIS values of the face prediction network on a trial-by-trial basis, support vector machines were used (Chang and Lin, 2011). Cross-validated decoding performance reached a maximum of 65.2% (mean performance 53.5%, SD 3.9% over subjects). When bonferroni correcting for the high number of subjects tested (n=52), for 22 of the subjects performance was still significantly better than for permuted datasets (p < 0.05/52). Note, that this is much more than would have been expected by chance (p = 1.1 x 10^-52^, binomial test).

### Analysis of information transfer

To understand how activated prior knowledge is communicated within the cortical hierarchy, we assessed the information transfer within the face prediction network in the prestimulus interval by estimating transfer entropy (TE, Schreiber, 2000) on source time courses for Face blocks and House blocks, respectively. Statistical analysis revealed significantly increased information transfer for Face blocks from aIT to FFA (p=0.0001, fdr correction) and from PPC to FFA (p = 0.0014, fdr correction). For House blocks information transfer was increased in comparison to Face blocks from brain area V1 to PPC (p=0.0014, fdr correction) (Figure 6).

**Figure 6.**
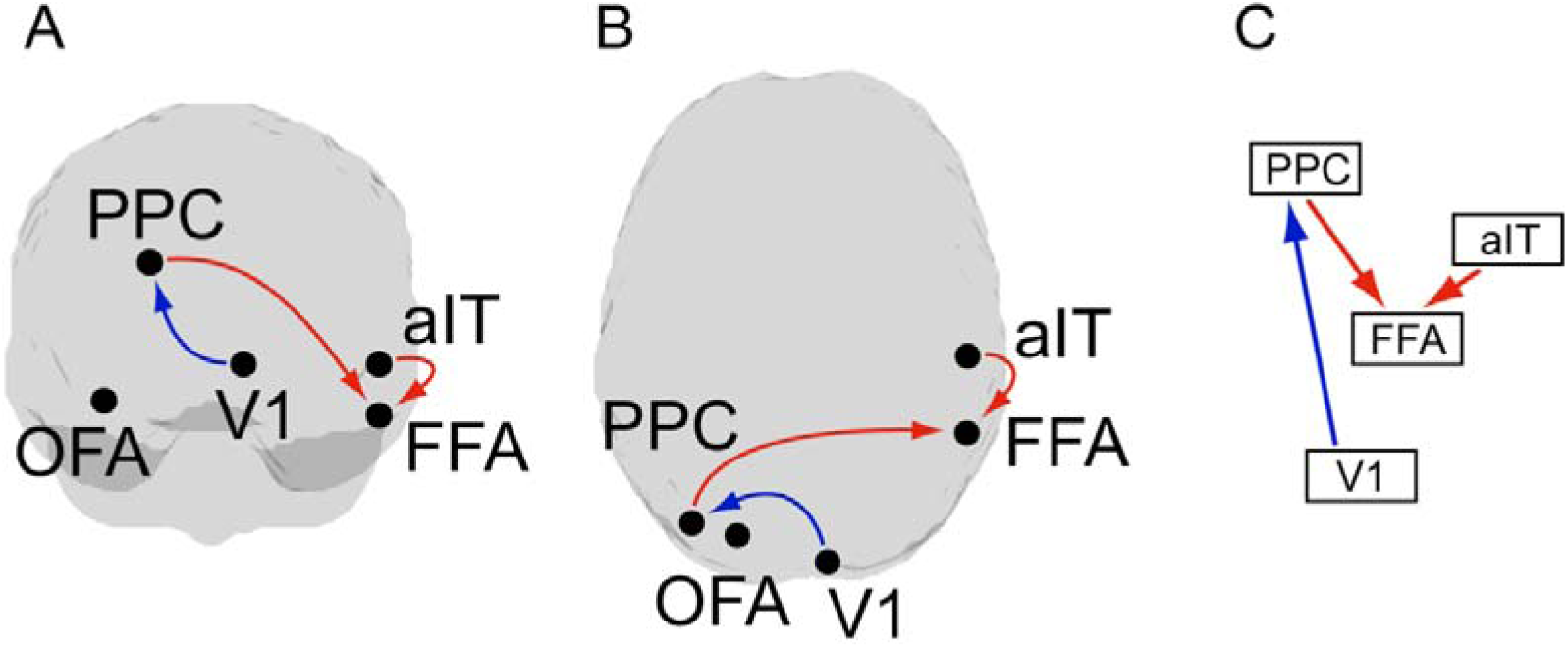
Analysis of information transfer in the prestimulus interval. Results of dependent sample permutation t-tests on transfer entropy (TE) values (Face blocks vs House blocks, n = 52, p< 0.05, fdr corrected). Red arrows indicate increased information transfer for Face blocks, blue arrows indicate increased information transfer for House blocks. Illustration of the resulting network in A) a view from the back of the brain, B) view from the top of the brain, C) depiction of the network hierarchy (based on the hierarchy in (Zhen et al., 2013; Michalareas et al., 2016).

### Post-hoc frequency resolved Granger causality

In order to investigate whether information transfer differences in Face and House blocks were reflected in specific frequency bands, we post-hoc performed a non-parametric spectral Granger causality analysis on the three links identified with transfer entropy analysis. For the link from PPC to FFA we found stronger Granger causality for Face blocks than House blocks in a cluster between 18 and 22 Hz (Figure 7, p=0.045, cluster correction for frequencies, uncorrected for the number of links in this post hoc test). The link from V1 to PPC showed a stronger Granger causal influence for House blocks than Face blocks between 94 and 98 Hz (Figure 7, p=0.042, cluster correction for frequencies, uncorrected for the number of links in this post hoc test). Using cluster correction, the link from aIT to FFA did not show significant differences in Granger causal influence.

**Figure 7.**
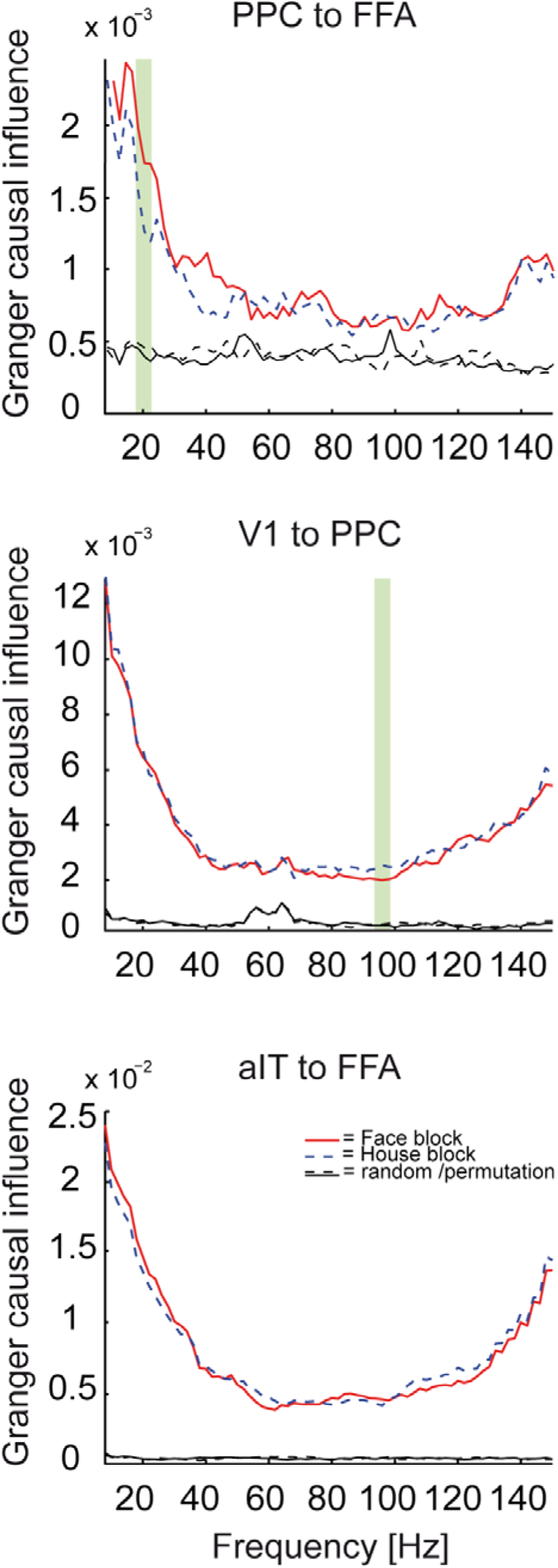
Frequency resolved Granger causality - post-hoc test for TE analysis. Granger causality for Face blocks and House blocks as well as random/permutation conditions. Green shaded regions indicate significant differences between Face and House blocks with cluster correction (dependent samples permutation t-test, n = 52, p<0.05). Frequency ranges were only considered as significant, if granger values for both block types in these frequencies also significantly differed from the random conditions (dependent samples permutation t-tests, n = 52, Bonferroni-Holms correction).

### Correlation of predictable information and reaction times

In order to study the association of predictable information and behaviour, we correlated the per subject difference of AIS values between Face blocks and House blocks with the per subject difference in reaction times. This analysis was performed for the three brain areas between which we found increased information transfer during Face blocks (FFA, aIT and PPC). For these brain areas we tested the hypothesis that predictable information for face blocks was associated with performance, i.e. reaction times during Face blocks. Negative correlation values were found for all of the three brain areas, however only brain area FFA reached significance when correcting for multiple comparisons: FFA robust Spearman’s rho −0.41, robust confidence interval (CI) after correcting for multiple comparisons [−0.68 −0.066]; aIT robust Spearmans rho = −0.12, CI [−0.4554 0.245]; PPC robust Spearman’s rho −0.21 CI [−0.5480 0.1178].

### Discussion

Here we tested the hypothesis that the neural correlates of prior knowledge activated for use as an internal prediction must show as predictable information in the neural signals carrying that activated prior knowledge. This hypothesis is based on the rationale that the content of activated prior knowledge must be maintained until the knowledge or the prediction derived from it is used. The fact that activated prior knowledge has a specific content then mandates that increases in predictable information should be found in brain areas specific to processing the respective content. This is indeed what we found when investigating the activation of prior knowledge about faces during face detection blocks. In these blocks predictable information was selectively enhanced in a network of well-known face processing areas. At these areas prediction content was decodable from the predictable information on a trial-by-trial basis and increased predictable information was related to improved task performance in brain area FFA. Given this established link between the activation of prior knowledge and predictable information we then tested current neurophysiological accounts of predictive coding suggesting that activated prior knowledge should be represented in deep cortical layers and at alpha or beta-band frequencies and should be communicated as a prediction along descending fiber pathways also in alpha/beta frequencies (Bastos et al., 2012). Indeed, within the network of brain areas related to activated prior knowledge of faces, information transfer was increased in top-down direction and related to Granger-causality in the beta band - in accordance with the theory.

We will next discuss our findings with respect to their implications for current theories of predictive coding.

#### 1. Activated prior knowledge for faces shows as predictable information in content specific areas

We found increased predictable information as reflected by increased AIS values in Face blocks in the prestimulus interval in FFA, OFA, aIT, PPC and V1. Out of these five brain areas FFA, OFA and aIT are well known to play a major role in face processing (Kanwisher et al., 1997; Kriegeskorte et al., 2007; Tsao et al., 2008; Pitcher et al., 2011).

In addition to increased predictable information in well-known face processing areas we also found increased predictable information in Face blocks in PPC. We consider the increase in predictable information in PPC also as content-specific, because regions in PPC have been recently linked to high-level visual processing of objects like faces (Pashkam and Xu, 2014) and activation of PPC has been repeatedly observed during the recognition of Mooney faces by us and others (Dolan et al., 1997; Grützner et al., 2010; Brodski et al., 2015).

In sum, our finding of increased predictable information for Face blocks in FFA, OFA, aIT and PPC confirms our hypothesis that activation of face prior knowledge elevates predictable information in content specific areas. Additionally, our results suggest that predictable information in content-specific areas is associated with the corresponding prediction on a trial-by-trial basis - by successfully decoding the anticipated category (Face or House block) from trial-by-trial AIS values at the face prediction areas.

However, while we found increased predictable information in content specific areas for Face blocks, we did not find brain areas showing increased predictable information for House blocks. Similarly, in a face/house discrimination task Summerfield and colleagues (2006b) observed increased activation in FFA, when a house was misperceived as a face. However, they failed to see increased activation in parahippocampal place area (PPA), a scene/house responsive region, when a face was misperceived as a house. The authors suggest that this might be related to the fact that PPA is less subject to top-down information than FFA – as faces have much more regularities potentially utilizable for top-down mechanisms than the natural scenes that PPA usually responds to. Additionally, because of their strong social relevance (e.g. Farah et al., 1995) faces capture attention disproportionally (e.g. Vuilleumier and Schwartz, 2001). Thus, also face predictions/templates may be prioritized in comparison to other templates e.g. for houses (Esterman and Yantis, 2009; Puri et al., 2009; Van Belle et al., 2010).

#### 2. Maintenance of activated prior knowledge about faces is reflected by increased alpha/beta power

We found a strong positive single-trial correlation of AIS with alpha/beta power for all face prediction areas. This finding supports the assumption that the maintenance of activated prior knowledge as indexed by AIS is related to alpha and beta frequencies.

Congruently with our findings, Mayer and colleagues (2015) recently showed that activation of prior knowledge about previously seen letters is associated with increased power in alpha frequencies in the prestimulus interval. Also, Sedley and colleagues (2016) observed that the update of predictions, which also requires access to maintained activated knowledge, is associated with increased power in beta frequencies.

Extending these previous findings, our results demonstrate that the activated prior knowledge usable as predictions for face detection is associated with neural activity in the alpha and beta frequency range (Bastos et al., 2012).

#### 3. Face predictions are transferred in a top-down manner and via beta frequencies

In Face blocks we observed increased information transfer to FFA from aIT as well as from PPC, both areas located higher in the processing hierarchy than FFA (e.g. Zhen et al., 2013; Michalareas et al., 2016). Thus, FFA seems to have the role of a convergence center to which information from higher cortical areas is transferred in order to prepare for rapid face detection.

Closely related to our findings (Esterman and Yantis, 2009) observed that anticipation effects for faces in FFA (and houses in PPA) were associated with increased activity in a posterior IPS region (part of the PPC) extending to the occipital junction. However, to our knowledge our study is the first to report face-related anticipatory top-down information transfer from PPC and aIT to FFA.

In addition to the two top-down links showing increased information transfer for Face blocks, we observed a bottom-up link from V1 to PPC with increased information transfer for House blocks. As we did not find a prediction network for houses and our analysis was thus only performed in the brain areas of the face prediction network, one can only speculate on the function of this bottom-up information transfer. It is possible that it indicates that house detection was rather performed in a bottom-up manner for instance by first identifying low level features that distinguish houses from their scrambled counterparts.

Our findings further demonstrated that information transfer in top-down direction was associated with Granger causality in the beta frequency band (PPC to FFA), while information transfer in bottom-up direction was associated with Granger causality in the high gamma frequency band (V1 to PPC).

The association of top-down information transfer with beta frequencies and bottom-up information transfer with gamma frequencies is in line with recent physiological findings in monkeys and humans (Bastos et al., 2015; Michalareas et al., 2016) and has been linked with predictive coding in Bastos’ microcircuit model (Bastos et al., 2012), resulting in the hypothesis of predictions being transferred top-down via low-frequency channels and prediction errors bottom-up via high frequency channels. In accord with this hypothesis, our group has recently shown that prediction errors are communicated in the high frequency gamma band (Brodski et al., 2015). Our present finding of top-down information transfer in low beta frequencies during anticipation of faces adds support to the microcircuit model hypothesis of a low-frequency channel for the top-down propagation of predictions (Bastos et al., 2012).

In line with our findings, the spectral dissociation between the transfer of predictions and of prediction errors recently received additional support from a MEG study applying Granger causality analysis for the investigation of information transfer during the prediction of causal events (Pelt et al., 2016). It should be noted that van Pelt and colleagues defined their network of interest for Granger causality analysis based on the prior assumption of the involvement of these brain areas in causal inference. In contrast, in our study defining the network via condition-specific AIS increases allowed finding the brain areas involved in predictive processing without relying on prior assumptions about their function.

#### 4. Pre-activation of prior knowledge about faces facilitates performance

Across subjects we found elevated predictable information in FFA in Face blocks in contrast to House blocks to be associated with shorter reaction times for Face blocks compared to House blocks. This suggests that especially pre-activation of prior knowledge about faces in FFA facilitates processing and speeds up face detection, as also suggested by FFA effects in previous fMRI studies (Esterman and Yantis, 2009; Puri et al., 2009). Our study is however the first to demonstrate that the size of the facilitatory effect on perceptual performance depends on the quantity of activated prior knowledge for faces in FFA, measurable as the difference in AIS between face and house block for each subject. Differential size of the faciliatory effect between subjects and the associated differences in the quantity of activated prior knowledge in FFA may be related to the differential ability in maintaining an object specific representation (see Ranganath et al., 2004).

## Acknowledgements

ABG received support by Ernst Ludwig Ehrlich Studienwerk (BMBF scholarship for graduate students). GFP received support by Villigst Studienwerk (BMBF scholarship for graduate students).

